# Sterilizing activity of spectinamide MBX-4888A when replacing linezolid in the Nix-TB regimen in the relapsing BALB/c mouse model of tuberculosis

**DOI:** 10.1101/2025.08.04.668403

**Authors:** Nathan Peroutka-Bigus, Michael S. Sherman, Firat Kaya, Samanthi L. Waidyarachchi, Jiuyu Liu, Joel Rushefsky, Michelle M. Butler, Terry Bowlin, Bernd Meibohm, Mercedes Gonzalez-Juarrero, Anne J. Lenaerts, Matthew Zimmerman, Richard E. Lee, Gregory T. Robertson

## Abstract

Spectinamides have garnered interest as experimental tuberculosis therapeutics owing to their safety profile and efficacy as partner agents when used in conjunction with established regimens in mice. The Nix-TB regimen of bedaquiline, pretomanid, and linezolid represents a short, effective regimen recommended for treatment of pre-extensively drug-resistant tuberculosis. However, linezolid administration is associated with severe adverse events which limits its use. Here we present preclinical data that spectinamide MBX-4888A can replace linezolid in Nix-TB.

## Main Text

The effective treatment of patients with all forms of tuberculosis (TB) is critical in the fight to eliminate the spread of drug-resistant *Mycobacterium tuberculosis* (*Mtb*). Combination antibiotic regimens intended for the treatment of patients with drug-resistant TB are associated with severe adverse events (AEs), which complicates treatment, leading to treatment discontinuation, non-compliance, and further expansion of drug resistance (1). The Nix-TB Phase 3 clinical trial demonstrated exceptional cure rates in patients with drug-resistant forms of TB using a three-drug regimen consisting of bedaquiline (B), pretomanid (Pa), and linezolid (L) (2). While the BPaL regimen is efficacious in treating drug resistant forms of TB, long-term administration of L was associated with AEs of peripheral neuropathy, myelosuppression, optic neuritis, and anemia, leading to dose reductions and frequent interruptions in L treatment (2). Spectinamide MBX-4888A (4888A) represents a new class of semisynthetic antibiotics that shows promising activity against *Mtb* in diverse preclinical murine models of TB disease with favorable safety profiles (3-9). Like L, 4888A targets protein synthesis, but occupies a different binding site in the ribosome (5).

Spectinamides are promising partner agents when administered with standard combination TB treatment regimens. The inclusion of spectinamide Lee-1599 with BPa resulted in increased bactericidal activity in a high dose aerosol subacute TB BALB/c infection model (4); while 4888A, given by injection with the front line standard of care regimen, led to improved bactericidal responses and treatment shortening in the subacute relapsing BALB/c model and in the more rigorous spectrum of disease chronic C3HeB/FeJ relapsing model of advanced pulmonary disease (8). An inhaled formulation of spectinamide Lee-1599 showed similar efficacy, but with reduced toxicities and an absence of known hematological effects, when used as a replacement for L in the BPaL regimen in BALB/c and C3HeB/FeJ mice (9). However, those studies failed to address whether a spectinamide when administered in place of L in BPaL resulted in durable sterilizing cure as measured by the prevention of relapse after treatment completion. This study employed the high-dose aerosol BALB/c relapsing mouse model to evaluate whether the injectable compound 4888A could replace L in the BPaL regimen, while still achieving a durable cure within a similarly short treatment duration.

One day following high dose aerosol with *Mtb* Erdman (see Methods in supplemental materials), average CFU lung burdens in female BALB/c mice were 4.04 log10 CFU and increased to 7.42 log10 CFU ten days later at the start of treatment. During treatment, both BPaL and BPa4888A significantly reduced *Mtb* lung burdens. With 4-weeks of therapy (5 of 7 days per week) BPa4888A reduced *Mtb* lung burdens by 4.98 log10 CFU, which was not significantly different from the 4.91 log10 CFU reduction with BPaL therapy (**Fig 1A, Table S1**). Following 8-weeks of therapy both treatments reduced *Mtb* lung burdens to below the limit of detection (> 6.54 log10 CFU), suggesting the presumptive clearance of *Mtb* from the lungs (**Fig. 1A, Table S1**). Weight loss is associated with severity of disease in this model. Following an initial period of weight decline, mice in both treatment arms gained weight throughout the remainder of the study (**Fig. S1**), indicating clinical improvements with treatment. Fewer relapse events were observed in mice held for 3 months after receiving 8-weeks of BPaL compared to those receiving 8-weeks of BPa4888A (29% vs 87% [P < 0.05]), with all mice achieving durable cure with 12-weeks of treatment (BPaL and BPa4888A) (**Fig. 1B**). Pharmacokinetic analysis based on sparse plasma sampling in infected (not shown) and healthy mice (**Fig. S2**) confirmed no drug-drug interactions with equivalent drug exposures for BPa and 4888A in plasma (**Table S2a-d**).

**Figure 1.**
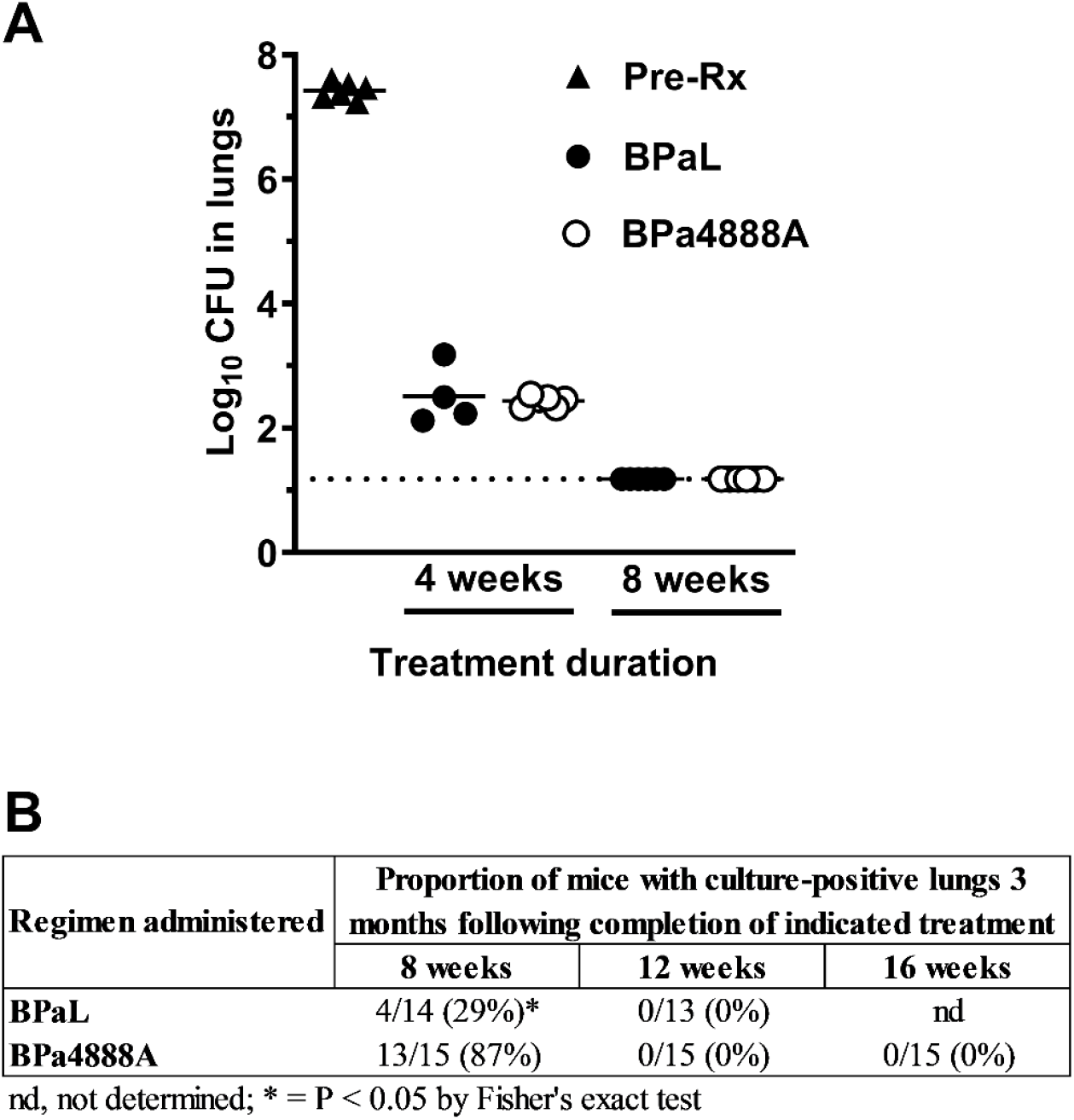
Antimicrobial regimen activity against *Mycobacterium tuberculosis* Erdman strain in BALB/c relapsing mouse model following high dose aerosol infection. **A**. Bactericidal activity of the two regimens over the 8-week treatment course. Pre-treatment (Pre-Rx) CFU lung burdens (closed triangles), 4-or 8-weeks of BPaL (closed circles), or 4-or 8-weeks of BPa4888A (open circles). **B**. The number of mice that relapsed 3 months after the indicated treatment duration over the group total. The percentage of mice relapsing is listed in parentheses. Drug doses in mg/kg: (B) Bedaquiline (25), (Pa) pretomanid (100), (L) linezolid (100), (4888A) MBX-4888A (200).

Although effective, clinical toxicities of L necessitate dose reduction or dosing holidays (2) which poses a serious threat to further expansion of clinical resistance to B (10). Modifications to BPaL, as the current WHO recommended pre-XDR therapy (11), are desirable to improve clinical control of TB. Although restricted currently to injection or inhaled therapies, spectinamides exhibit favorable safety profiles and tolerability in preclinical studies (3, 8, 9), and as shown here, appear capable of contributing to durable cure in the BALB/c relapsing mouse model as a replacement for L in the BPaL regimen. Indeed, in a previous study, 4888A was administered subcutaneously for 20 weeks in *Mtb*-infected mice, providing an additional data point regarding the safety and tolerability of spectinamide 4888A (8). Our studies further the available data on the utility of spectinamides in the treatment of TB. While these results are presently limited to outcomes in the conventional BALB/c relapsing mouse model - thought to best represent uncomplicated TB disease - our prior work demonstrated effective distribution and accumulation of 4888A within the caseous lesions of C3HeB/FeJ mice (8), suggesting a possible role for BPa4888A in harder to treat patients with necrotic acellular caseum filled granulomas; this remains to be tested.

This study has several limitations. BPa was not tested as a direct comparator, so the contribution of 4888A to treatment efficacy and cure cannot be estimated. However, previous work showed that a closely related spectinamide, Lee-1599, does add appreciably to the BPa backbone in this same infection model (4). Further studies are needed to de-convolute the contribution of 4888A to cure in this preclinical model. Additionally, while sparse PK assessment provided data that co-administration of 4888A with BPa did not result in any drug-drug interactions, a full PK analysis was not performed, and the comparative behaviors of L and 4888A in this context were not evaluated.

The fact that both BPaL and BPa4888A achieved a 100% cure rate after 12 weeks of treatment, and that 4888A was well tolerated, adds further evidence supporting the safety and efficacy of this antibiotic. Given its unique binding site and absence of current clinical use, it is anticipated that 4888A will not be impacted by any pre-existing clinical resistance – which is not the case for linezolid (a repurposed drug) for which clinical resistance is on the rise (12, 13). One counterpoint was a significant early advantage of BPaL in reducing the number of mice that relapsed at the 8-week timepoint. As both combination regimens were anchored by BPa, one possible explanation is the nearly 100% oral bioavailability of L in humans and mice (14, 15) improves early target attainment and drives cure in drug-tolerant populations that arise during drug treatment, but other possibilities such as absolute target potency and the differences in L (16) and 4888A (17) model-based exposure responses should also be considered.

## Acknowledgements

This research was supported by the National Institute of Allergy and Infectious Diseases and the Office of the Director of the National Institutes of Health (grants R01AI090810 [R.E.L.], R44AI098271 [Microbiotix]). The content is solely the responsibility of the authors and does not necessarily represent the official views of the National Institutes of Health. We acknowledge the staff of the Laboratory Animal Resources at Colorado State University for animal care. Bedaquiline fumarate was obtained through the NIH HIV Reagent Program.

